# Social hierarchy is established and maintained with distinct acts of aggression in male *Drosophila*

**DOI:** 10.1101/2020.05.12.091553

**Authors:** Jasper C. Simon, Ulrike Heberlein

## Abstract

Social interactions pivot on an animal’s experiences, internal states, and feedback from others. This complexity drives the need for precise descriptions of behavior to dissect the fine detail of its genetic and neural circuit bases. In laboratory assays, male *Drosophila melanogaster* reliably exhibit aggression, and its extent is generally measured by scoring lunges, a feature of aggression in which one male quickly thrusts onto his opponent. Here, we introduce an explicit approach to identify both the onset and reversals in hierarchical status among opponents and observe that distinct aggressive acts reproducibly precede, concur, or follow the establishment of dominance. We find that lunges are insufficient for establishing dominance. Rather, lunges appear to reflect the dominant state of a male and help in maintaining his social status. Lastly, we characterize the recurring and escalating structure of aggression that emerges through subsequent reversals in dominance. Collectively, this work provides a framework for studying the complexity of agonistic interactions in male flies enabling its neurogenetic basis to be understood with precision.

## INTRODUCTION

Agonistic interactions essential for establishing social hierarchies over food, territory, or mates often progress as a sequence of stereotyped acts. Like other goal-driven behaviors [1], the steps along these sequences are triggered by context and modified by past experience [2,3], through largely unknown mechanisms that are coupled to an animal’s internal state [4]. Determining these mechanisms remains difficult, however, because social exchanges proceed as a dynamic, reciprocal, and continual feedback loop of interactions between two (or more) decision-making individuals that progresses over both moment-by-moment and protracted times scales [5]. Inter-male adversarial contests in the fruit fly *Drosophila melanogaster* are well suited for studying such complex social phenotypes, due to their established study in a laboratory setting [6,7] and the multitude of techniques available for manipulating and recording gene and neuron function [8]. Nonetheless, efforts to explain which behaviors lead to dominance hierarchies [9,10] have suffered from a lack of consensus on methods and results for studies of aggression (as discussed within [11,12]).

Here we use an explicit criterion to characterize the temporal relationship among various aggressive acts and the establishment and reversals in dominant hierarchical status between pairs of male flies in arenas commonly used for high-throughput phenotyping [13,14]. Using this approach, we observed that the acts of fencing, boxing, and tussling appear well positioned in time to be related to the establishment of social dominance, whereas lunging and chasing appear related to its maintenance. We verified that lunging, the most frequently used behavior for quantifying aggression, unexpectedly occurs nearly exclusively after the onset of dominance and is therefore unlikely to be involved in its establishment. In fights between high- versus low-lunging genetic lines, males became dominant at chance levels. This result suggests that the total number of lunges executed by an individual is insufficient for explaining his social outcome. We further observed that lunging persists as dominant males continue to combat familiar opponents, and also when they confront unfamiliar opponents in novel settings. Lastly, we report that a fraction of the aggressive acts surveyed intensified through subsequent reversals in social dominance, adding to a fuller appreciation of this complex, recurrent social exchange. Together, this work provides a framework for untangling which aggressive acts causally relate to the establishment of social hierarchy and those that are a consequence.

## RESULTS

### Identifying hierarchical relationships

Upon introduction to experimental arenas, pairs of males display independent and haphazard movements that typically includes one or both males jumping, taking flight, and/or climbing the wall as they attempt to escape the arena. With time, the males settle into a period of exploration during which they eventually meet. This initial contact ends their independent movements and leads to bouts of short, alternating pursuits that are characteristically restricted to the floor, which was entirely composed of food (see **Methods**). This engagement then predictably progresses to a situation wherein one male consistently pursues the other until the retreating male leaves the floor, presumably in an attempt to flee by climbing the wall. It is this diagnostic “pursue-to-climb” sequence that we use as criterion to identify the initial establishment of a hierarchical relationship, with the pursuing male being classified as dominant (**Figure 1** and **Movie 1**). This diagnostic criterion is also used to identify reversals in social status, as discussed later within this work.

**Figure 1.**
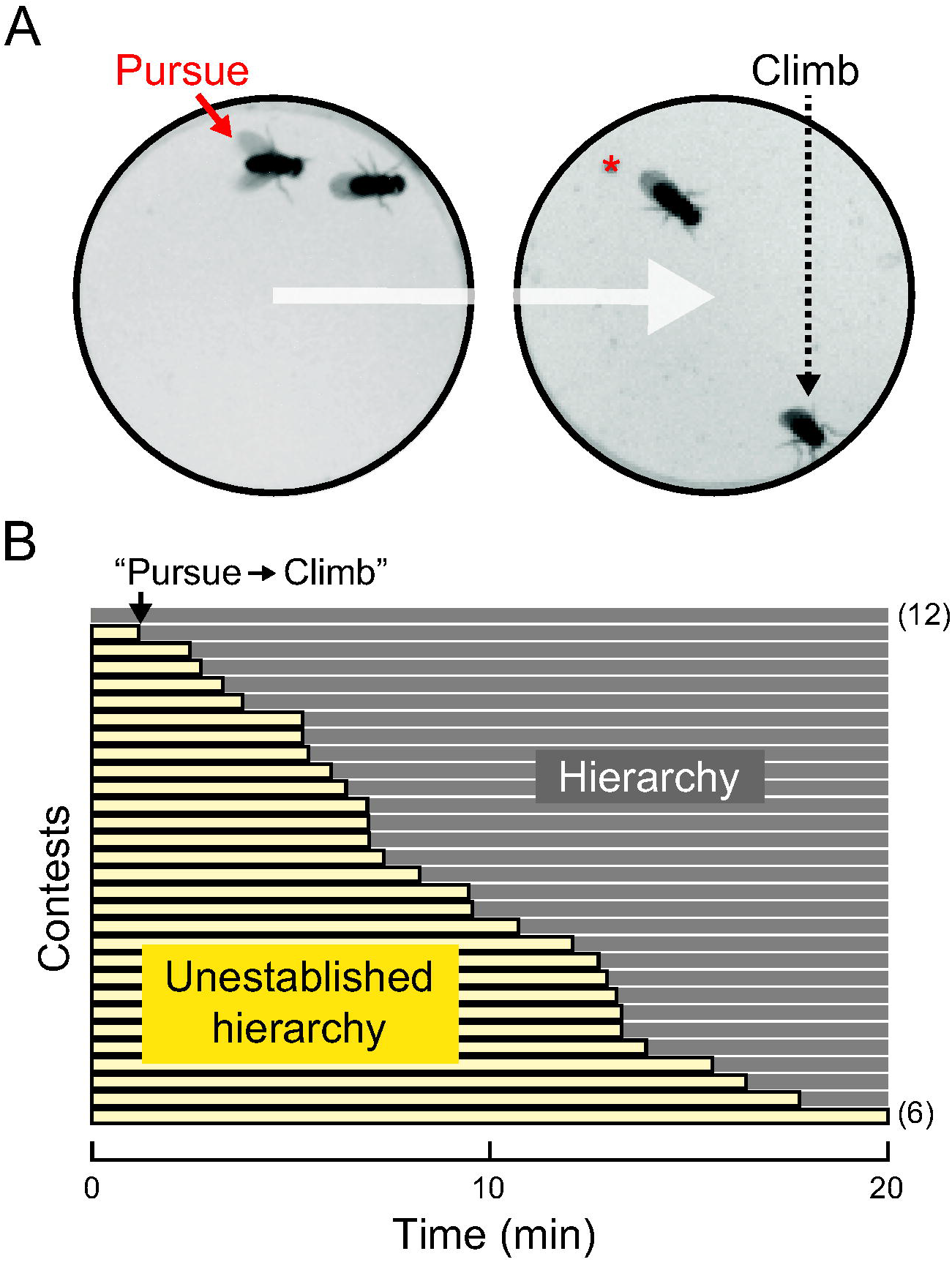
Identifying the onset of social hierarchy within arenas used for studies of aggression. (A) Example movie images illustrating the diagnostic “pursue-to-climb” criterion for identifying the onset of social hierarchy. Pursuing males (red, solid arrow), that often proceed to guarding the floor (red, asterisk), are classified as dominant and those then attempting to flee by climbing the wall of the arena as subordinate (black, dashed arrow). (B) Twenty-minute contests ordered as rows from early to late onset of establishment (top to bottom). Using the diagnostic criterion, clear hierarchies form in the majority of pairings. Periods of unestablished social standing (light yellow bars) precede clear social hierarchy (gray bars). Contests in which hierarchies formed prior to observation (full gray bar, top row, n=12) and those in which hierarchies never formed (full yellow bar, bottom row, n=6) did not change results and were excluded from further analysis.

### Distinct aggressive acts precede, concur, or follow the establishment of dominance

We observed adversarial contests in experimental arenas that have been widely used to study aggression [13,14]. These arenas, which have a floor comprised entirely of food, increase the frequency of flies’ interactions by providing constant access to a food resource and yet retaining an uncluttered background with no food cup and no decapitated female [15]; moreover, the entirety of the chamber is within view of the camera. This arrangement permits use of computer vision methods to help automatically annotate various aggressive acts over time [13], followed by direct manual inspection for correcting erroneous annotations and identifying the onset of dominance. Together this complementary workflow allows for a robust, high-resolution portrayal of the aggressive acts displayed during the progression of agonistic interactions.

Males who had never experienced a fight (hereafter, “naïve males”) were introduced together as pairs into arenas. This simultaneous introduction lessened the disparity in which individuals discovered the contested resource and allowed uninterrupted observation of their interactions leading up to the onset of social dominance. Over a period of 20 minutes, we measured the occurrence of several acts, including wing flicks and the following acts reported to span the range of aggression, listed here from low to high intensity: fencing, chasing, lunging, boxing, and tussling [6]. Note that chasing is a long, high-speed trailing behavior that is distinct from pursuits, which are shorter. At times we found boxing and tussling difficult to distinguish, and thus for ease of scoring they were combined into the single category, “Box, Tussle,” as done by others [6]. See **Figure S1** and **Table S1** for details of behavioral classification, annotation, and software used.

From both individual contests and aggregate data, we observed reproducible patterns of behavior (**Figure 2**). Males consistently displayed the highest levels of “Fence,” from the beginning of contests until the onset of dominance. Thereafter, we observed only sporadic displays of “Fence” with no clear temporal structure. In contrast, “Box, Tussle,” appeared abruptly, peaked, and then sharply decreased immediately prior to the onset of dominance (**Figure 2D**), and on occasion also transiently reappeared preceding subsequent reversals in social status. “Wing flick” was observed broadly, first occurring with low frequency at the beginning of contests and then making a salient uptick near the onset of dominance, with males continuing this elevated level of “Wing flick” throughout the remainder of the contest. Unexpectedly, “Lunge” was rarely observed before the establishment of dominance, but began and continually increased in intensity thereafter. Occurrences of “Chase” were never observed prior to dominance and emerged well after its establishment (see analyses associated with **Table S2**; as reported previously [6]). Together, by aligning recordings from individual contests to the onset of social dominance, we have exposed which acts may causally relate to its establishment and those apparently more related to its maintenance.

**Figure 2.**
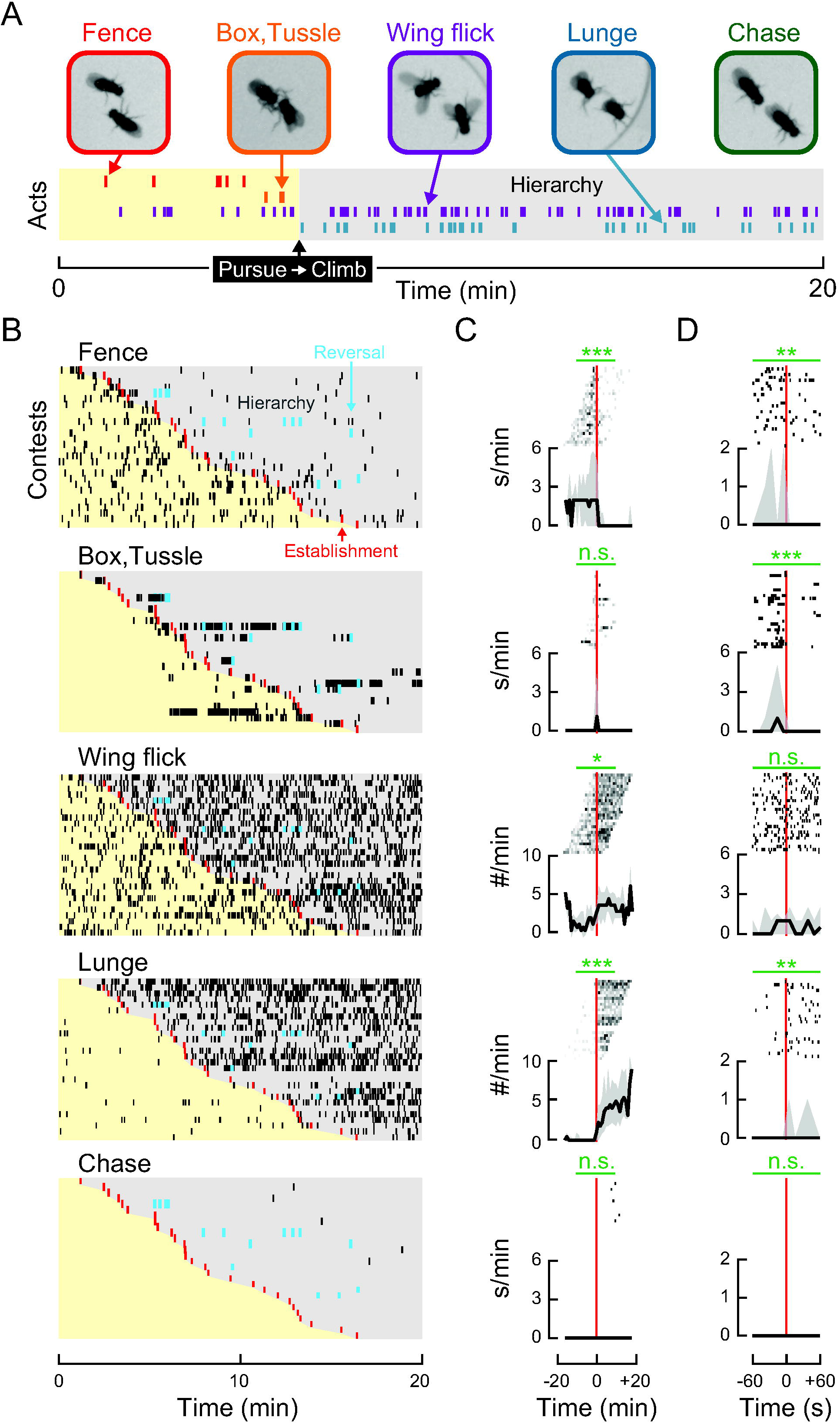
Fence, box, and tussle interactions precede the establishment of social hierarchy, whereas the majority of wing flicks, lunges, and chases follow. (A) Raster plot from a single contest illustrates the relationship between the observed reproducible sequence of behaviors and the establishment of social hierarchy. Each row and color corresponds to a different act. (B) Raster plots of examined behaviors displayed from early to late in sequence (top to bottom) show the temporal structure of discrete acts (black ticks). Within each plot, individual contests between pairs of naïve males are ordered as rows by latency to the onset of establishment (red ticks). After establishment males exhibit clear hierarchical relationships (gray shading), with occasional reversals in dominance (blue ticks). (C, D) Corresponding peri-event plots aligned to the onset of establishment (vertical, red line) with acts displayed above associated collective medians (black line) and interquartile ranges (gray envelope). (C) Entire contests with acts binned into one-minute intervals. (D) ±one-minute windows with discrete acts. (C, D) Statistical comparisons for the frequency of acts within five-(C) and one-minute (D) windows are noted above plots (horizontal, green lines). In all cases the Wilcoxon signed rank test was used to assess whether differences in the observed acts exist between paired, equal-length windows of time before and after the establishment of social hierarchy. See **Figure S2** for details regarding data exclusion. Lunging events in figure were automatically classified and are displayed without correction (see **Methods**). Common nomenclature for statistical significance used throughout study: not significant (n.s.), *p*<*.05* (*), *p*<*.01* (**), *p*<*.001* (***), *p*<*.0001* (****), *p*<*.00001* (*****).

### Lunges are insufficient for explaining hierarchical outcomes

The act of lunging is commonly used for studying aggression (see reviews [11,16]), with several studies suggesting they play a critical role in establishing dominance [9,17,18]. We, however, observed lunging almost always after the establishment of dominance (as seen in **Figure 2**). To confirm this finding, we ran a second, independent group of contests and reviewed the incidences of lunging by direct inspection with slow playback to correct for false negative and semi-automatically for false positives displays (see **Methods**). After careful review, it was clear that males nearly exclusively lunged after the establishment of dominance, with almost all of the lunges executed by the dominant male (**Figure S3**).

From the repeated observation that lunging occurs after the onset of dominance, we reasoned that lunges were unlikely to play a role in its establishment. If this were true, then the number of lunges executed by a male over the 20-minute contest would be unrelated to whether an individual emerges dominant. To test this, we set up contests with mixed pairs of males from previously identified genotypically high- and low-lunging lines (see **Methods**) and measured the total number of lunges executed by those males that emerged dominant. Males from genotypically high- and low-lunging lines became dominant at chance levels regardless of their opponents’ genotypes (**Figure 3A**). Moreover, we saw no influence of the total number of lunges on dominance outcomes, even when males from mixed pairs executed dramatically lopsided amounts of lunging (see fourth column in **Figure 3B**; **Figure 3 – Supplement 1** for additional examples and

**Figure 3.**
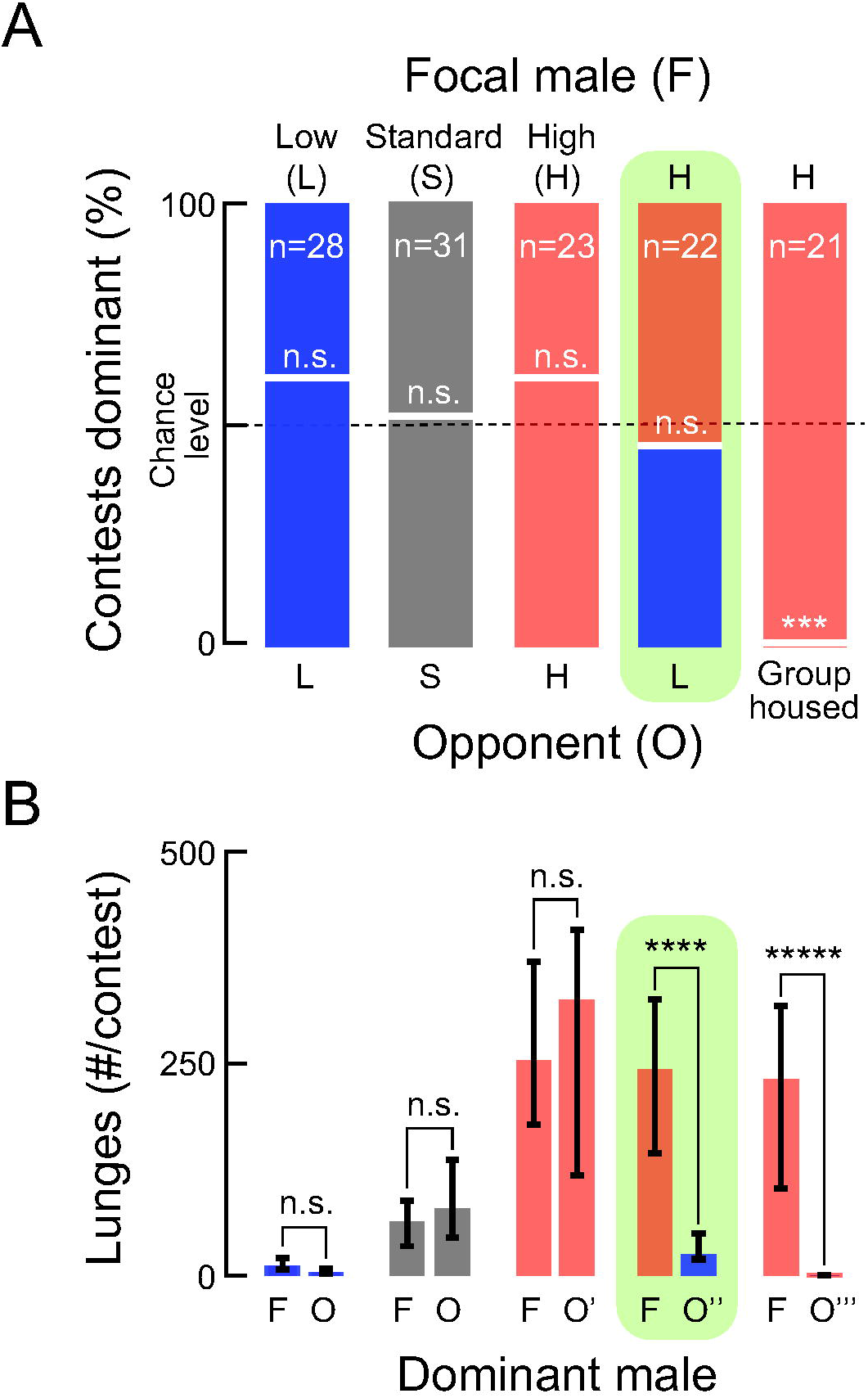
Number of lunges executed by dominant individuals is insufficient for explaining hierarchical outcomes. (A) Focal males (F, above) and opponents (O, below) were paired and the percentage of contests in which males became dominant are shown within each stacked bar chart. Focal males from genotypically low (L, blue), standard (S, gray), and high (H, light red) lunging lines became dominant at chance levels when paired with same-genotype opponents. For the tested pairing between mixed high (H, light red) and low (L, blue) lunging males also became dominant in comparable amounts (shaded in green). In contrast, for contests in which high-lunging focal males (H, light red) were paired with socialized opponents (group housed) from the same genotype, focal males became dominant 100% of the time (compared to chance levels; Fisher exact, *p*=*.0005*; right-most bar). In all cases males were socially isolated unless noted. (B) Medians with interquartile ranges for the total number of lunges executed by dominant individuals from the contests reported in “A” above. The number of lunges executed by dominant focal males (F) from low (blue), standard (gray), and high (light red) lunging lines were no different than when those of paired opponents (O) from the same genotypes became dominant. Focal males from a high-lunging line (F, light red) displayed a comparable, high level of lunging when paired with all opponents (from left to right): males from the same genotype (O’, light red), genetically low-lunging males (O’’, blue), or socialized males (O’’’, light red) from the same genotype (Kruskal-Wallis, H(3)=5.92, *p*=*.1142*). However, in tested pairings the levels of lunges executed by the dominant, high-lunging focal males (F, light red) were significantly greater than those of dominant, low-lunging opponents (O’’, blue; Wilcoxon Rank-Sum, Z=3.9244, *p*=*.00009*; shaded in green) and socialized opponents from the same high-lunging line (O’’’, light red; Wilcoxon Sign, *p*<*.00001*). The designation of “Focal” is independent of which male’s wing was clipped for identification (see **Methods**), with the sole purpose of clarifying experiments.

**Table S2** for details regarding genetic lines). It was therefore unsurprising to observe that males from both the high- and low-lunging genotypes executed the majority of lunges following the establishment of dominance (**Figure S4**). Taken together, the results that males nearly exclusively lunge after the establishment of dominance and that the number of lunges executed by dominant males is unpredictive of social outcomes provide evidence that lunges do not establish dominance, and more likely play a role in its maintenance.

### Males express dominance by lunging at both familiar and unfamiliar opponents

For the majority of contests, we observed that only a single, stable hierarchical relationship formed between pairs of naïve males (as shown in **Figure 2** and **Figure S3**). One in five contests, however, included subsequent reversals in social status after the initial establishment of dominance, with lunges, sometimes by both males, clustering near each reversal. To better understand how these lunges relate to reversals in status, we verified which individual executed each lunge (see **Methods**). As in contests with only a single establishment of dominance, males nearly exclusively lunged after the initial establishment of dominance. Thereafter, currently and previously dominant males displayed the majority of lunges preceding reversals, and only dominant males lunged following the final reversal in status (**Figure S3**). This analysis provides evidence that lunging is a good indicator of dominance status, and supports the interpretation that through subsequent reversals in dominance lunges play a role in maintaining a male’s social status (also see shaded, green box **Figure S5C-E**).

In the experiments described above, both increased total time holding dominance and also the occurrence of reversals in social status contributed to a persistent high level of lunging by individuals against familiar opponents. Moreover, it has been reported that “winner” males lunged and became dominant earlier in contests against naïve opponents [19,20]; and also, that males appeared to change their fighting tactics as a consequence of winning or losing [21]. Therefore, to test if lunging functions as a general tactic for maintaining dominance irrespective of opponent or location, we paired the emergent dominant males from a first contest between two naïve males (**Figure 4A**) with naïve males in unexplored arenas (**Figure 4B**). The most salient feature of these second contests was that the majority of previously-dominant males lunged prior to establishing dominance (**Figure 4B, C**), although their paired naïve opponents did not. This pattern of behaviors was unlike that of pairs of naïve males, which rarely lunged before the establishment of dominance (**Figure 4A, C**). The temporal sequence for all other aggressive acts studied was comparable to those observed in contests between two naïve males (data not shown). Finally, to examine if the precocious lunging helped previously-dominant males assert dominance against unfamiliar males in unexplored arenas, we quantified the number of contests in which the previously-dominant males lunged prior to asserting dominance. For the majority of these contests, previously-dominant males successfully asserted their dominance (**Figure 4D**). Taken together, these results support the understanding of lunging as a general expression of dominance that aids in maintaining a male’s social status.

**Figure 4.**
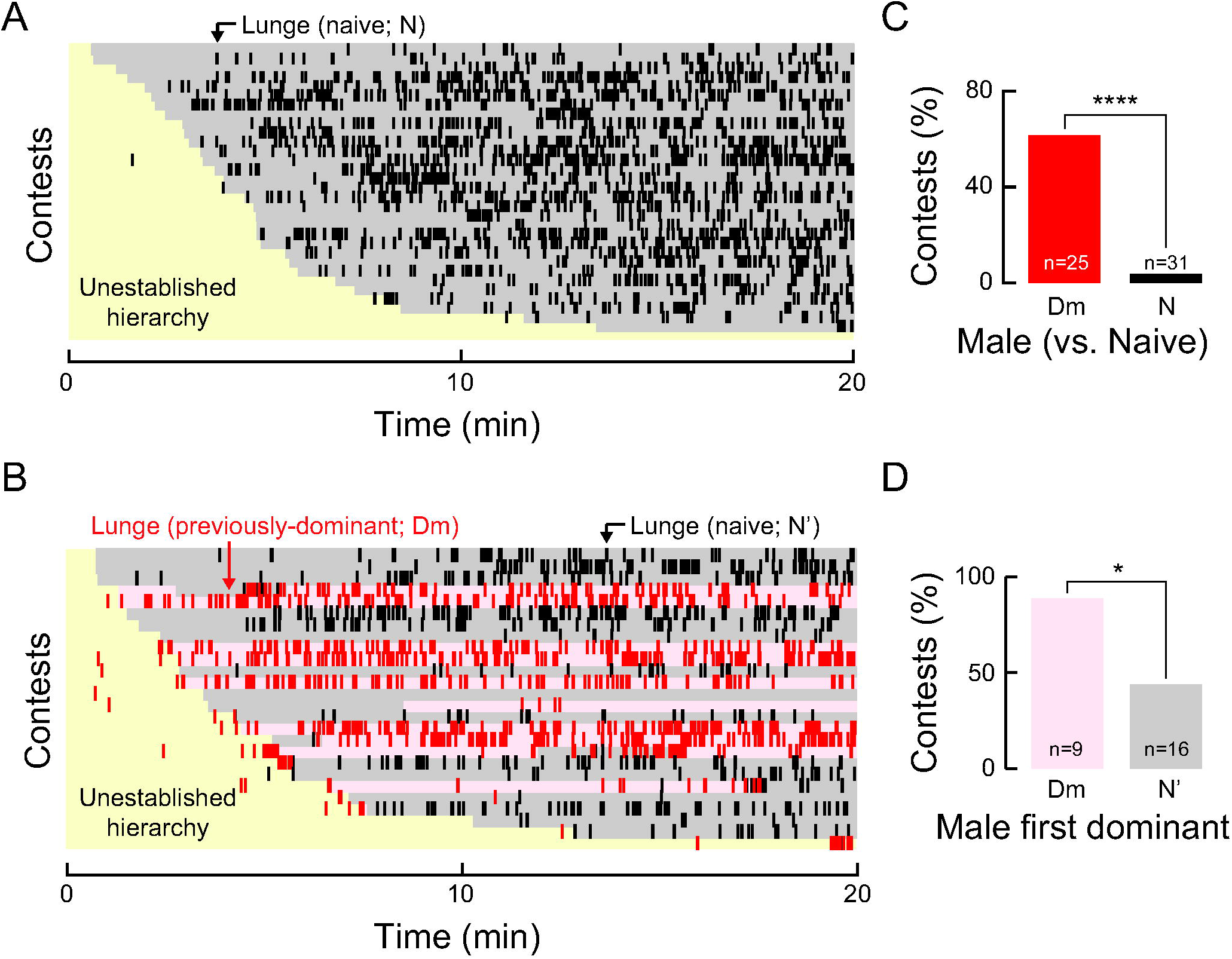
Lunging persists as previously-dominant males confront unfamiliar opponents in unexplored arenas. (A, B) Raster plots show the temporal structure of lunges executed by naïve (black ticks) and previously-dominant (red ticks) males in relation to establishing social hierarchy. Within each plot, individual contests between pairs of males are ordered as rows by latency to the onset of establishment. Periods preceding establishment (yellow shading), and the prevailing dominance status for naïve (gray shading) and previously-dominant (light red shading) males are noted as observed within 20-minute contests. (A) Pairs of naïve males. (B) Previously-dominant males paired with naïve opponents in a second, unexplored arena. (C) The number of contests in which the previously-dominant males (red bar; “Dm”) lunged prior to asserting dominance over naïve males was significantly greater than those in which randomly selected individuals from pairs of naïve males (black bar; “N”) lunged prior to establishing dominance (Chi-Square: *X*^2^ (1, N=56) = 16.07, *p*=*.00006*). (D) The number of contests in which previously-dominant males (light red bar; “Dm”) lunged prior to asserting dominant was also significantly greater than those in which they lunged early, failed, and became subordinate to naïve males (gray bar; “N’”; Chi-Square: *X*^2^ (1, N=25) = 4.89, *p*=*.02701*).

## DISCUSSION

Social dominance forms when individuals yield to the agonistic advances of others, often in the context of conflict over resources such as access to food, territories, and reproductive opportunities [22]. The relationship between specific aggressive acts and the establishment of social hierarchies remains contentious largely for they appear so highly correlated, even with disagreements over which drives the other [23]. In this work, we use an explicit approach to characterize with high temporal resolution how a range of known acts of aggression relate to the establishment, maintenance, and subsequent reversals in dominant hierarchical status.

The “pursue-to-climb” criterion introduced within this work has several advantages. It is easy to describe and straightforward to implement. It accommodates that adversaries compete over resources and includes an operationally defined ‘escape,’ both of which are important ethological considerations [6,24]. Further, it provides the means to align and thus compare the onset of dominance, which varies in time across contests due to differences in various influences (e.g., inherited factors, past social histories, current internal states, handling by the experimenter, environmental conditions, and feedback between adversaries). Lastly, it can be used to update the classification of dominance if and when reversals in status occur after an initial hierarchical relationship has been established.

A significant challenge to studying complex social behaviors such as those related to the establishment of social hierarchy is that the numerous interactions possibly regulating them are often highly correlated with outcomes. By using a criterion to identify the establishment of dominance that is independent from the particular behaviors studied, we avoid a circular definition of social hierarchy, and thus allow for stronger claims of causality between specific aggressive acts and its emergence (for a graphical summary **Figure 5**).

**Figure 5.**
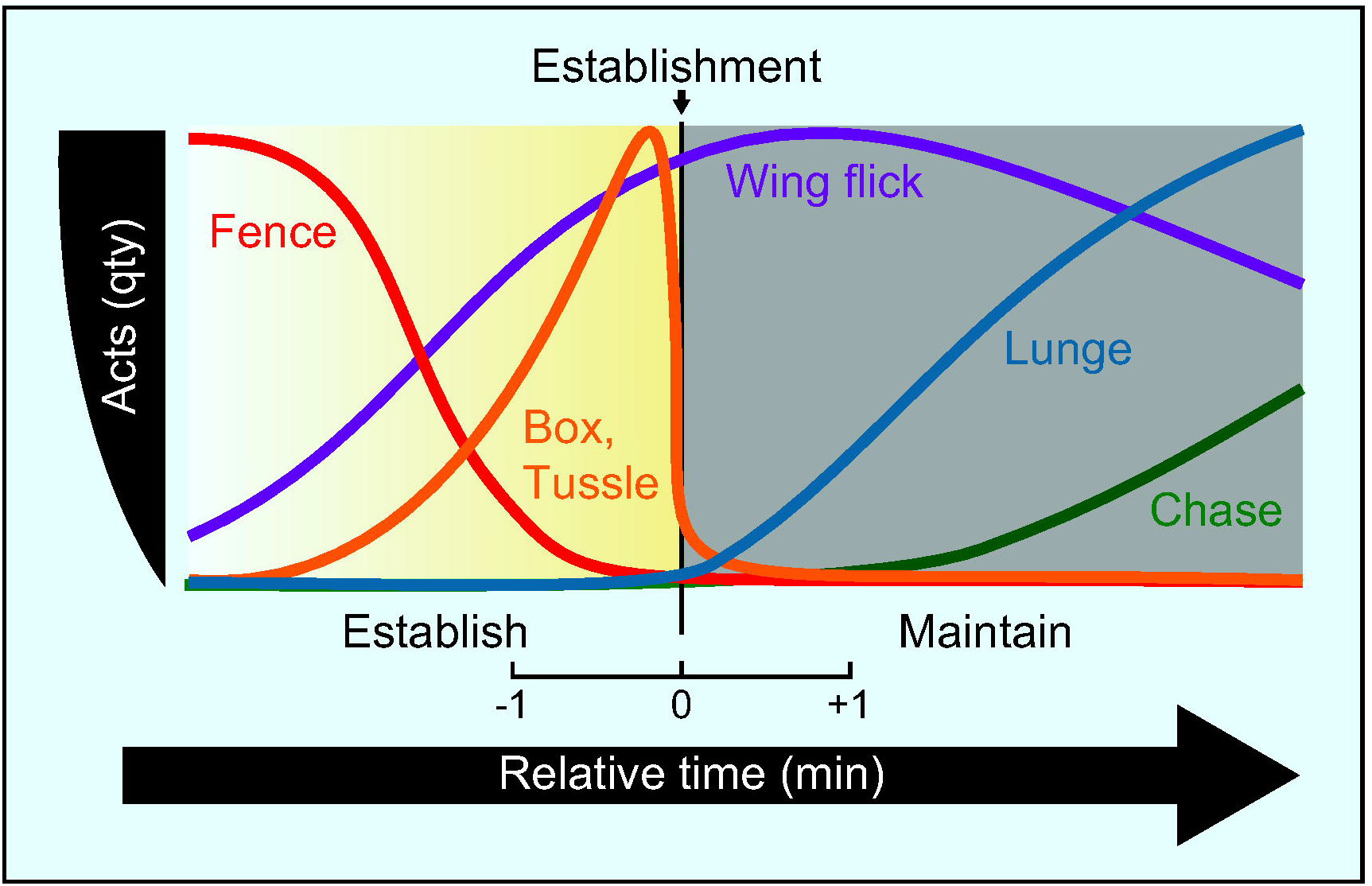
Sequence of aggressive acts during the establishment of social hierarchy. After adjusting to the experimental arena, pairs of males typically recognize each other as opponents. Successively they then exhibit fencing (red line) and then display low levels of wing flicks (purple line). As contests escalate (graded yellow box) fencing bouts decrease in frequency and are replaced by boxing and tussling (orange line) concomitantly with an increasing number of wing flicks. Transiently the amount of boxing and tussling peaks and then sharply drops immediately preceding the establishment of dominance (time = 0). Lunging (blue line) follows the establishment of dominance, is the principal aggressive act during the maintenance of dominance (gray box), and is accompanied by a steady display of wing flicks. Finally, increased chasing (green line) observed after some time suggest stabilized hierarchical outcomes.

Unexpectedly, we show that almost always only once naïve males establish (or have held) dominance do they lunge, and moreover they continue lunging against familiar opponents even after a dominance hierarchy is settled (as has been described previously [9,25]). From this and our observation that previously-dominant males precociously lunge and successfully assert dominance over unfamiliar males in novel settings, we propose that lunging is likely uninvolved in the establishment of dominance, but rather symptomatic of a changed internal state [26] and plays a role in maintaining an individual’s social status.

The complexity of reversals in dominance requires more study. In an attempt to more fully understand the role that the examined aggressive acts played in the reversal of hierarchical status, we analyzed a larger number of contests that included reversals. From a preliminary characterization of these contests, lunges appeared as just one of several interacting acts likely involved in this process (**Figure S5**). It is clear, as with the initial establishment of dominance, that further consideration is needed to formally conclude which acts of aggression causally drive these dynamic, reversing, and escalating exchanges.

In summary, we demonstrate that identifying biological meaningful epochs of time with a criterion that is independent of precise descriptions of behavior, helps in establishing both nuanced and holistic understandings of this complex behavioral phenotype. These understandings should assist in refining the roles of known acts of aggression [6,27,28], and possibly discovering new ones. Finally, when used in conjunction with high-throughput screens, the approach and work described herein provide a framework to clarify the genetic and neural mechanisms underlying the coordination of behaviors involved in establishing hierarchical social relationships in insects.

---

**Movie 1. Established hierarchical relationship.** Example movie from a 20-minute contest showing an established hierarchical relationship, wherein a subordinate male repeatedly attempts to climb the wall of the arena while a dominant male (clipped wing) guards the floor.

## SUPPLEMENTAL FIGURES

**Figure S1.**
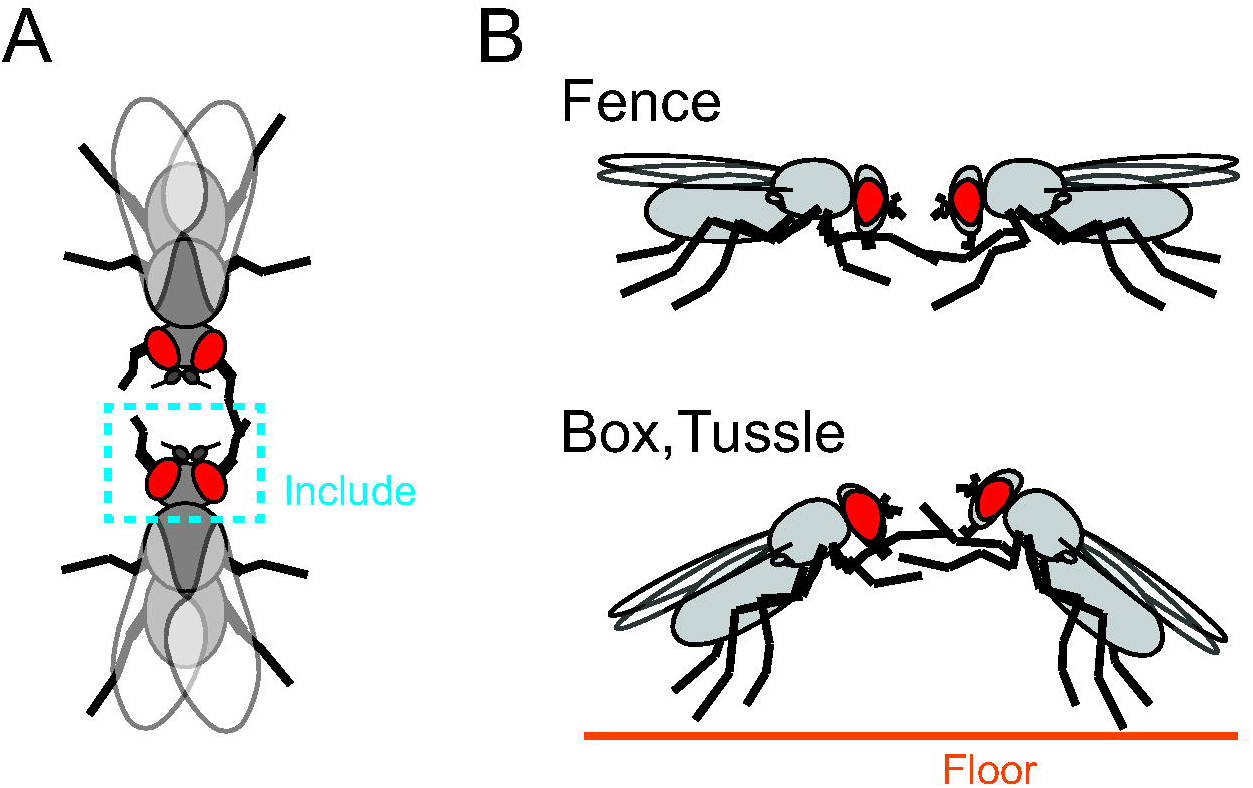
Details for including and classifying “Fence” versus “Box, Tussle” interactions. (A) Illustration drawn from a top viewing angle to clarify the criteria for inclusion. For an interaction to be included the focal male must touch the front legs or head of the opponent (blue dashed box). (B) Side views showing the postural difference between “Fence” and “Box, Tussle.” Interactions that included touch from a prone stance were classified as “Fence.” Any touching when one or both males exhibited an erect posture with both front legs lifted off the floor were classified as “Box, Tussle.” Interactions in which males disengaged, directly executed a 360-degree rotational turn, and then reengaged were annotated as a single continuous act.

**Figure S2.**
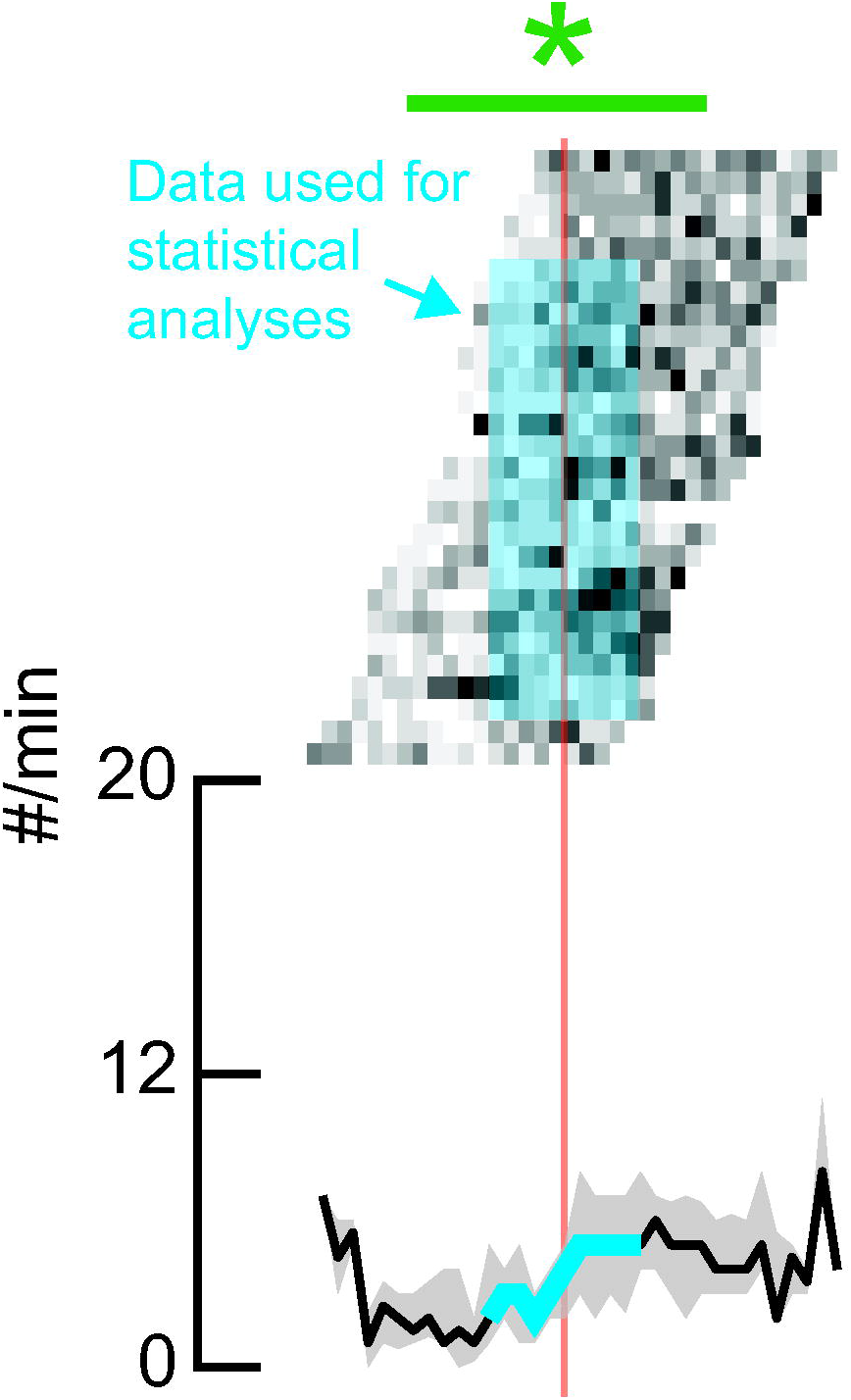
Example data to show the requisite exclusions for balanced statistical analyses. An example dataset shows the data used for statistical analyses. In order to apply paired statistical tests, only contests allowing five minutes or more (always equal) windows of time before and after the establishment of social hierarchy were used unless otherwise noted. In the example shown only contests with five minutes of recorded data before and after establishment of dominance were used (blue shaded box, above; median frequency of behavior for corresponding period denoted as bold, blue line, below). In this example, data recorded earlier than five minutes preceding and following establishment and also the data from the entire first five and last two contests were excluded. The exclusions decreased the sample size, and thereby the statistical significance, yet never qualitatively changed results.

**Figure S3.**
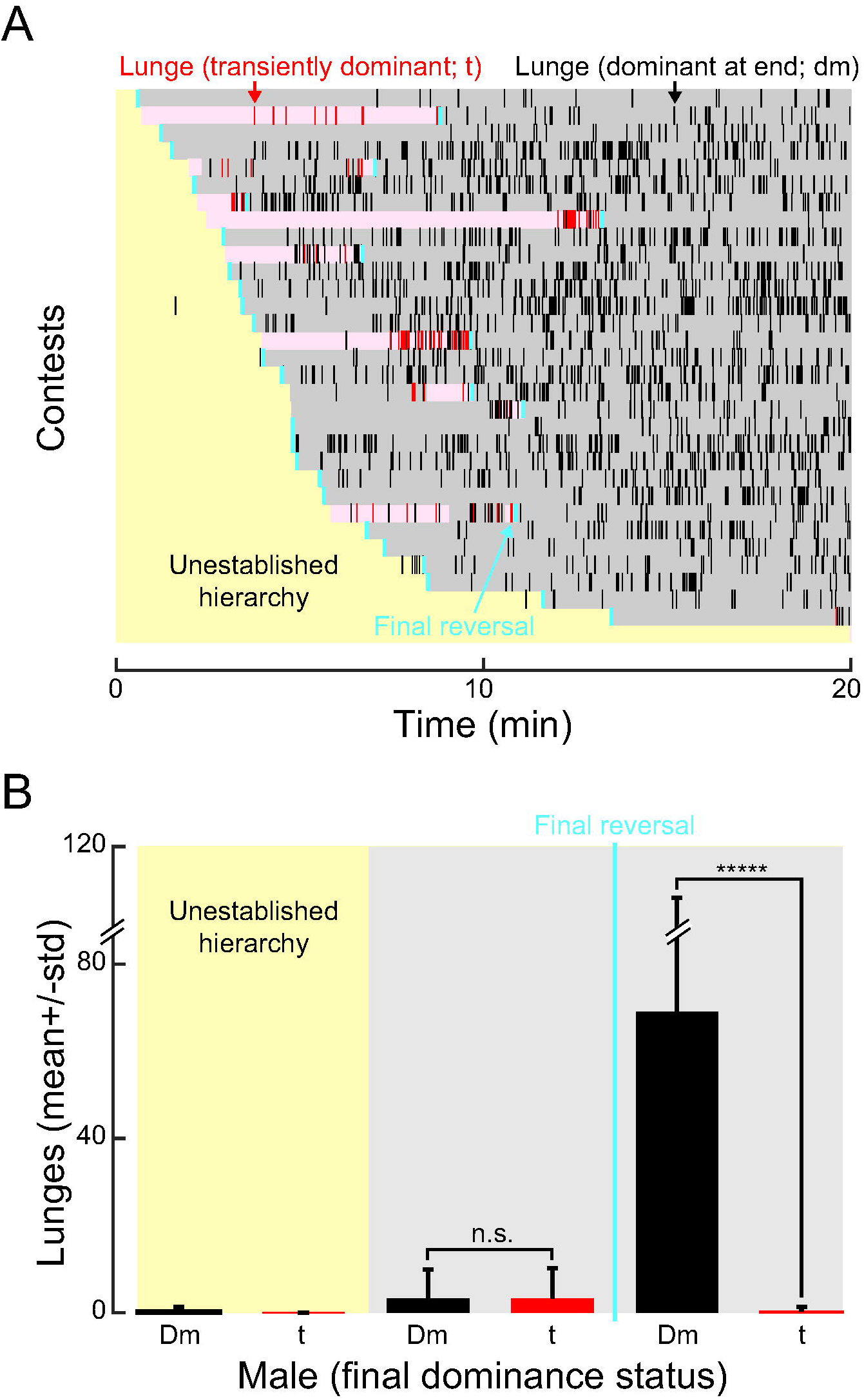
Males express dominance by lunging. (A) Raster plot displaying the temporal structure of lunges executed by transiently dominant males (red ticks; t) and those observed dominant at end (black ticks; dm) in relation to the initial establishment and reversals in social dominance. Individual contests between pairs of naïve males are ordered as rows by latency to the onset of establishment. Periods preceding establishment (yellow shading), ultimately transient (light red shading), and of final dominance status (gray shading) are noted as observed within 20-minute contests. Final reversals in social status for each contest are indicated with blue ticks (ticks for intermediate reversals were excluded for clarity). (B) Lunging reflects dominance. Uncommonly, lunges were observed before the initial establishment of dominance (bar plots highlighted by yellow shading). In the period following the initial establishment yet before final reversals in dominance (vertical, blue line), currently and previously dominant males executed the majority of the lunges in comparable, low amounts irrespective of final dominance outcomes. Thereafter, all lunges came from dominant males (Dm, black bar), and subordinate males (t, red bar) displayed none (Wilcoxon rank sum, Z=7.1057, *p*<*.00001*). This figure includes the identities of which male lunged and also denotes any reversals in social status between the pairs of naïve males as observed within the contests reported in **Figure 4A.**

**Figure 3 – Supplement 1.**
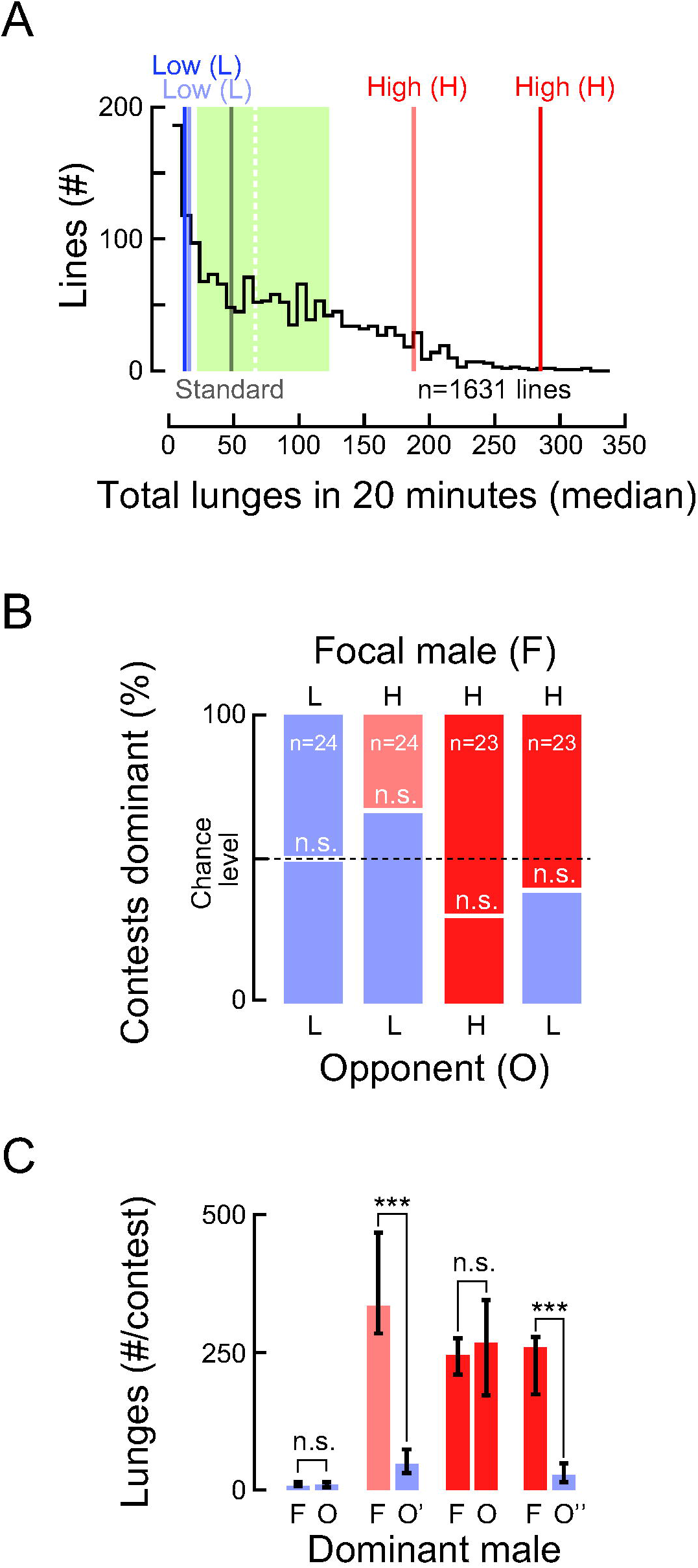
Independently isolated genotypically low, standard, and high lunging lines identified from a P-element screen; additional genotypically high- and low-lunging lines corroborate findings in Figure 3. (A) The distribution of P-element lines screened before outcrossing is skewed right (black, stair-step line) with the median (vertical, white dashed line) and interquartile range (green shading) centered around 67 total lunges, as calculated by summing the total lunges executed by pairs of same-genotype males from 20-minute contests. The median number of lunges executed by the genotypic standard (gray) used throughout the study and also the low (blue, light blue) and high (light red, red) lines are shown after outcross. (B) Focal males (F, above) and opponents (O, below) were paired in various combinations and the percentage of contests in which males became dominant are shown within each stacked bar chart, with adversaries coded by letter and color as indicated in “A” above. Focal males became dominant at chance levels in all contests. (C) Lunging executed by dominant males in both same- and mixed-genotype pairings were consistent with those reported in **Figure 3**. Dominant focal or opponent males arising from same-genotype pairs displayed equivalent amounts of lunges (F vs. O, light blue and red). Focal males from an additional high-lunging line (F, red) displayed a comparable, high level of lunging regardless of opponent (Kruskal-Wallis test; H (1)=0.0039, *p*=*0.9503*). However, in tested pairings the levels of lunges executed by both the original and the additional dominant, high-lunging focal males (F, light red; F, red) were significantly greater than an additional dominant, low-lunging opponent (vs. O’, light blue; Wilcoxon Rank-Sum, Z=3.8884, *p*=*.00010* and versus O’’, light blue; Wilcoxon Rank-Sum, Z=3.8111, *p*=*.00014*).

**Figure S4.**
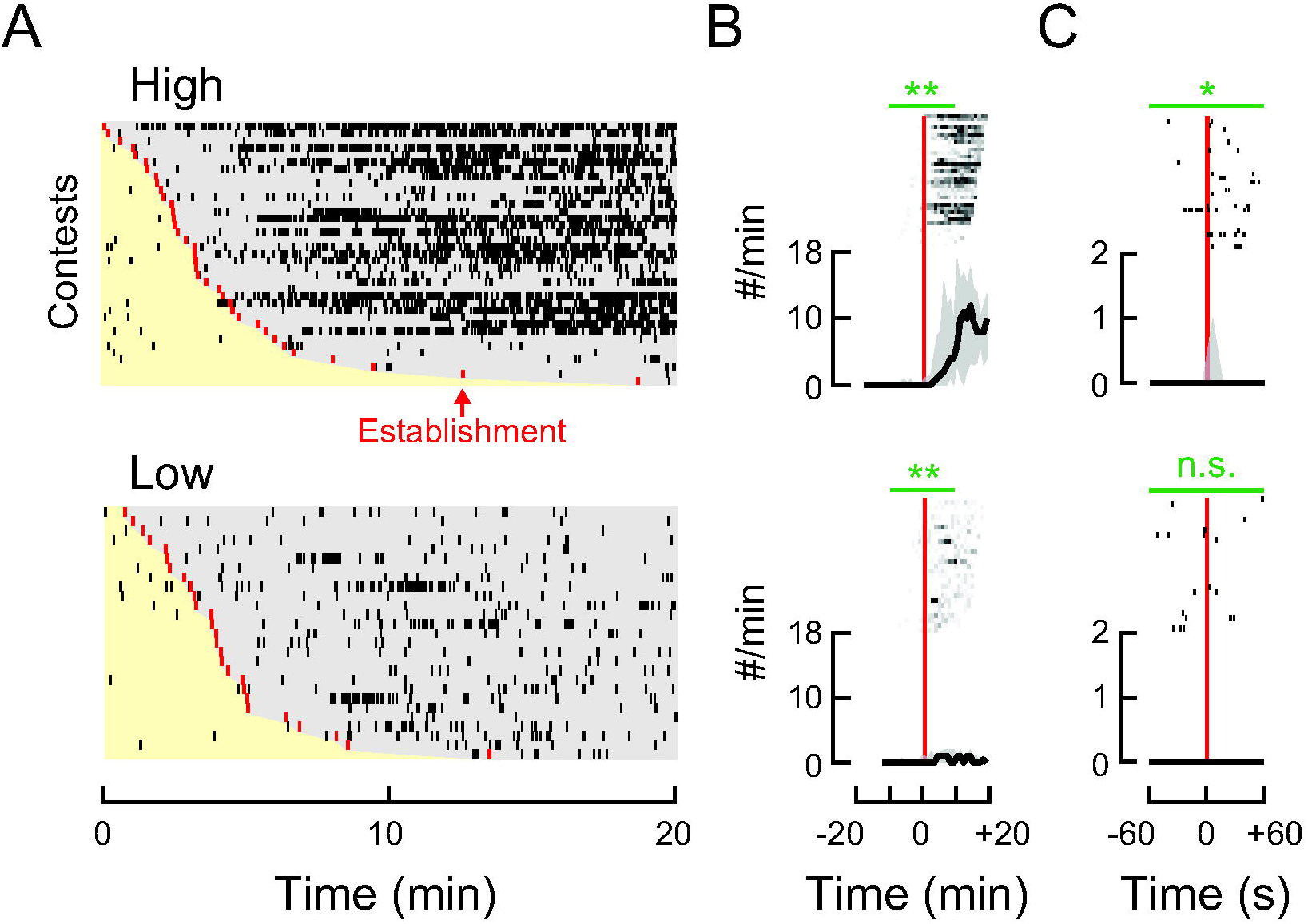
Lunges were executed after establishing dominance irrespective of genotype. (A) Raster plots show the temporal structure for lunges executed by males from genotypically high (top) and low (bottom) lunging lines (lunge = black tick). Within each plot, individual contests between pairs of naïve males from the same genotype (high vs. high and low vs. low) are ordered as rows by latency to the onset of establishment (red ticks). After the establishment of dominance males exhibit clear hierarchical relationships (gray shading). (B, C) Corresponding peri-event plots aligned to the onset of establishment (vertical, red line) with lunges displayed above associated collective medians (black line) and interquartile ranges (gray envelope). (B) Entire contests with lunges from each contest binned into one-minute intervals. (C) ±one-minute windows with distinct lunging events. (B, C) Statistical comparisons for the frequency of lunges within five- (B) and one-minute (C) windows prior to and following establishment are noted above plots (horizontal, green lines). In all cases the Wilcoxon signed rank test was used as described in **Figure S2**. Lunging events in figure were automatically classified and are displayed without correction (see **Methods**).

**Figure S5.**
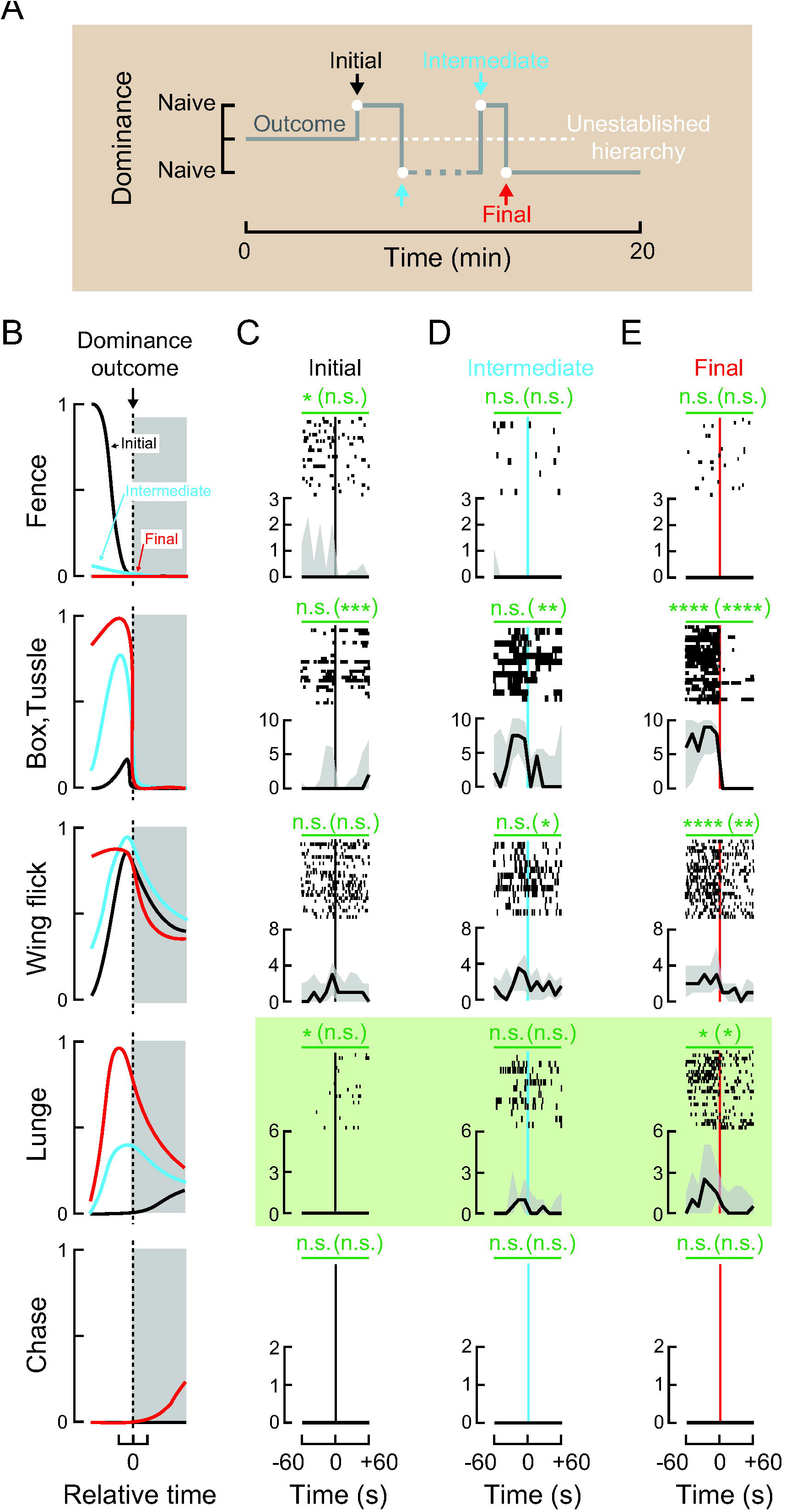
Recurring aggressive acts intensify through subsequent reversals in dominance. (A) Schematic illustrating the establishment and subsequent reversals in dominant hierarchical status in contests between pairs of naïve males. The gray, switching line indicates the state of dominance with colored arrows indicating the onset (white dot) for the initial establishment (Initial, black), subsequent reversals (Intermediate, blue), and the final hierarchical outcome (Final, red). (B) Idealized results for the observed changes of aggressive acts among the initial establishment (black line) and intermediate (blue line) and final (red line) reversals of dominance status (gray shaded box). A value of “1” indicates the highest level for any particular act to help clarify its relative increase or decrease; drawings are based on the data in **Figure 2**, those reported here in “C-E,” and contests using males from the genotypically high- and low-lunging lines (data not shown). Depicted in the top panel of “B,” early on males exhibit the highest levels of “Fence,” which then drops during the minute immediately preceding the initial establishment (black line). From then on only intermittent and low levels of fencing were observed through subsequent reversals (Intermediate, blue and Final, red lines). In contrast, as shown in the next panel down, the amount of “Box, Tussle” peaks during the minute immediately preceding the initial establishment (black line), and then peaks again and again, increasing and broadening through subsequent reversals (Intermediate, blue and Final, red lines). (C-E) Peri-event plots with ±one-minute windows aligned to the initial establishment (C; vertical, black line), the intermediate subsequent reversals (D; vertical, blue line), and final outcome (E; vertical, red line) of social hierarchy. Discrete acts are displayed above associated collective medians (black line) and interquartile ranges (gray envelope). (C-E) Statistical comparisons for the frequency of aggressive acts within ±one-minute and ±10-second (within parentheses) windows prior to and following establishment are noted above plots (horizontal, green lines). In all cases the Wilcoxon signed rank test was used as described in **Figure S2**. Lunges increased, formed clusters, and the peaks of clusters shifted forward in time relative to each subsequent reversal (C-E; shaded, green box). Lunging increased in a scalable manner (One-way ANOVA: df(2), F(22.5470), *p*<*.00001* and followed by post hoc comparisons: Initial versus Intermediate, *p*=*.0135*, Intermediate versus Final, p=*.0475*, and Initial versus Final, *p*<*.00001*; all with Bonferroni correction). The shift in peaks was quantified as a procession index; PI = (Lunges *minute preceding* – Lunges *minute following*) / (Lunges *minute preceding* + Lunges *minute following*). One-way ANOVA: df(2), F(6.1421), *p*=*.0049* and followed by post hoc comparisons: Initial versus Intermediate, *p*=*n.s.*, Intermediate versus Final, *p*=*n.s.*, and Initial versus Final, *p*=*.0046*; all with Bonferroni correction). The intensity for several of the aggressive acts increased alongside that of lunges. Collectively, the characterization exposes a recurring structure that emerges during protracted engagements – one in which both males reciprocally escalate fights just prior to reversals in status – seemingly in attempt to maintain or reclaim social dominance. Data within this figure came from contests that all contained reversals in dominance (n=25) identified from larger set of adversarial contests (n=146).

## SUPPLEMENTAL TABLES

**Table S1.** Details of the aggressive acts reported within manuscript. Descriptions and process for generating the data used to characterize the temporal structure of the various aggressive acts reported within manuscript.

| Act | Description | Method | Annotation | Software |
| --- | --- | --- | --- | --- |
| Fence | Extending one or both forelegs and pushing against the forelegs or head of opponent from a prone position. | Manual | **Figure S1.** | VCode [29] |
| Box, Tussle | Rearing up on hindlegs while striking, pushing, or pulling opponent with forelegs. | Manual | **Figure S1.** | VCode [29] |
| Wing flick | Brief spreading followed by furling of one or both wings. | Automatic | Left and right wings flicks from both focal and opponent males outputted by CADABRA were combined (using Matlab) as follows:<br>wingFlick =<br>sort([wing.fli.r.obj1.t<br>wing.fli.l.obj1.t<br>wing.fli.r.obj2.t<br>wing.fli.l.obj2.t]) | CADABRA [13] |
| Lunge | Forward thrust of body with outstretched forelegs while snapping down on opponent. | Semi-Automatic | Assigning identification to time-stamped lunges classified by CADABRA (see Supplementary Tables 2,3 therein for details) provided information for which male executed a particular lunge. | CADABRA [13]<br>followed by<br>scoreMoviesLunges [20] |
| Chase | Running after opponent. | Automatic | First male following another at a speed greater than 5mm/s for $\geq 1$s (see Supplementary Table 9 within [13] for details). | CADABRA [13] |

**Table S2.** Behavioral measures for the out-crossed P-element lines used within manuscript. P-element lines ordered by row from fewest to greatest number of lunges (median ±quartile) per 20-minute contest as identified within screen after outcrossing. Line names indicate batch followed by trial number and come from the labelling scheme of our laboratory. Color and intensity may be used to associate lines with results from **Figure 3** and **Figure 3 – Supplement 1**. In addition to its high levels of lunges, the line 9.3 was chosen for its high amounts of chase (see below). Consequently, to support the notion that chase events follow the establishment of social hierarchy as observed in **Figure 2**, we analyzed chase events in contests between pairs of males from 9.3 and also in mixed contests between males from 9.3 and the those from the low-lunging line 11.261. In both cases, we observed that all chase events occurred following the establishment of hierarchy (compared to equal numbers of chase events before and after establishment; 9.3 versus 9.3, Fisher exact, n=18; *p*=*.0010*; 9.3 versus 11.261, Fisher exact, n=11; *p*=*.0351*). Together, combining the above results with those from **Figure 2**, we report that we have only observed chases following the establishment social hierarchies, Fisher exact, n=34, *p*<*.00001*.

| Line | Color | Intensity | Sample size<br>(n=contests) | Tussle | Wing flick | Lunge | Chase |
| --- | --- | --- | --- | --- | --- | --- | --- |
| 11.261 | Blue | Low | 62 | 1 (0/4.3) | 66.5<br>(12/128) | 12.5<br>(1/26) | 0 (0/1) |
| 15.46 | Light blue | Low | 64 | 1 (0/4) | 66<br>(24.5/213.8) | 16<br>(6.5/37.5) | 0 (0/1) |
| 5.116 | Gray | Standard | 301 | 3 (0/8) | 95 (57/149) | 48<br>(23.5/104) | 0 (0/3) |
| 9.3 | Light red | High | 24 | 2 (0.9/7.6) | 138<br>(73/212.6) | 184<br>(98.5/236) | 4<br>(0/16) |
| 11.27 | Red | High | 52 | 3 (1/8.3) | 138<br>(85.9/214.9) | 277<br>(186/391) | 0 (0/2) |

## MATERIALS AND METHODS

### Fly lines

All *Drosophila* lines originate from a P-element collection generated by the Heberlein laboratory that were then outcrossed for at least six generations to *w*^*1118*^ Berlin.

### Animal rearing

To lessen developmental heterogeneity, males used in experiments were collected from lines maintained with controlled densities (by seeding vials with 5 males crossed to 10 females and removing these parental animals after 3 days), reared in customary 8-dram plastic vials on standard media (cornmeal/yeast/molasses/agar), maintained at 25°C and 65% relative humidity, and entrained on a 12:12-hour light-dark cycle. The lights-on phase started at 1pm EST. Transitions between dark and light were immediate.

### Animal handling

To model a uniform, probable ‘ecological baseline’ amount of social experience including to have had mated, males for experiments were collected after 7 hours, yet within the first 24 hours following eclosion using CO_2_. For experiments in which the identity of individuals was required, during collection, a small section of wing chosen randomly from one of an eventual pair was removed, a procedure shown to be inconsequential to the outcomes of adversarial contests [20].

### Animal housing

In order to increase or suppress the level of aggression, males were housed individually in 10-mm diameter × 75-mm tall glass tubes (Fisher, Waltham, MA) or as groups of 15 males in 8-dram vials [30,31]. For both housing conditions flies had *ad libitum* access to standard media and were incubated in the same conditions as they were reared.

### Animal observation

Experiments were performed on 4~6-day old males that were removed from food immediately prior to start. Trials began after a 30-minute adjustment period to “lights on” and the environmental conditions of the observation room. All runs were completed within the first 4 hours of “day.” Replicate experimental and control trials were intercalated and duplicate batches of trials were run over several days at least twice, weeks to months apart.

### Aggression assay

Unless noted, aggression assays were performed similarly to those previously described [26] modified from [14]. Briefly, pairs of males were aspirated simultaneously into a 16-mm diameter × 10-mm tall enclosed arena staged in an environmentally-control room held at 25°C and 45% humidity and their behavior was recorded for 20 minutes. To encourage consistent aggressive interactions, the entire floor of the arena was composed of a thin layer of apple juice-sugar agar made as 10g sucrose and 9g agar boiled in 400mL 100% apple juice (Mott’s, Plano, TX). To keep the quality of the agar floor consistent, it was used either immediately after a 2-hour setting period or air-dried for 1 hour, wrapped inside of plastic (Saran, Racine, WI), and held at room temperature until the following day. To impede flies from climbing, the wall of the arena was made slippery by coating it with Fluon (BioQuip, Rancho Dominguez, CA). Similarly, to limit flies from hanging from the ceiling, the lid of the arena was brushed with Sigmacote (Sigma-Aldrich, St. Louis, MO), a transparent silicone paint allowing an unobstructed view for recording from above. Both coatings were left to dry for at least 24 hours before running experiments.

### Aggression screen

The genetically high- and low-lunging males used came from a P-element screen (Mark Eddison, J.C.S, U.H.; more details will be published elsewhere; see **Figure 3 – Supplement 1A** for the distribution of median total lunges for all lines observed in screen). From this screen, only lines that maintained a stable phenotype after outcrossing, had normative levels of activity as estimated by measuring their total distance travelled and total number of jumps, and appeared otherwise healthy were used. A line, 5.116, exhibiting normative levels of aggression and activity was also identified in the screen. This “standard” was used for all principal experiments reported within this work. **Table S2** includes measures of the aggressive acts from the various lines used within the manuscript.

### Data acquisition

Interactions between males were recorded at 30 frames per second using a Basler A622f camera (Edmund Optics, Barrington, NJ), with movies from an array of arenas saved simultaneously with gVision video acquisition software [32] for further analysis. Experimental arenas were lighted from behind with a flicker-free, uniform, white backlight (Coherent, Santa Clara, CA), making recordings suitable for machine vision tracking and behavioral classification methods. For the principal experiments reported in this work, recordings started within 2 minutes after introduction. For experiments analyzed from the P-element screen, recordings started 5-7 minutes after introductions, allowing sufficient time for loading and 5 minutes for flies to settle.

### Behavior classification

To catalog the changes in social hierarchy and also the aggressive acts, we used a combination of manual and automatic classification methods (**Figure S1** and **Table S1** for methods, descriptions, and software used). In summary, the establishment and reversals of social dominance and also the aggressive acts, fence, box, and tussle, were manually annotated with VCode [29] and we used CADABRA [13] to automatically classify wing flick, lunge, and chase. To assign specific lunges to identified males we used a semi-automated method previously reported [20]. This software application uses as inputs both original full-length movies and corresponding time-stamped behavioral events (generated with CADABRA [13] in our case) and iteratively makes short movies inclusive of each event, allowing users to accurately assess if or which male executed a particular lunge. For the P-element screen, pairs of males were tracked and automatically scored for total number of wing flicks, tussles, lunges, chases, and jumps using CADABRA [13].

### Statistical analysis

Data processing, plotting, and statistical analyses were all conducted in Matlab (MathWorks, Natick, MA). Details for analyses are explained and reported within each figure legend. See **Figure S2** for specifics regarding data exclusion.

## Code availability

Documentation and code used for verifying the identification of which male lunged is available at: https://github.com/JasperCSimon/scoresMovie.git.

## Acknowledgments

We thank Erik Hoopfer for gifting us his experimental rig and sharing code to run CADABRA [13] on the Janelia computer cluster; Mehrab Modi, Lisha Shao, Mark Eddison, and Kit Longden for insightful suggestions; and past members of Heberlein laboratory for feedback and support. This work was funded by HHMI.

## Authors Contributions

J.C.S. proposed project, designed, conducted, analyzed experiments, and wrote the manuscript; U.H. supported, advised project, and helped to write manuscript.

